# White matter microstructure in women with acute and remitted anorexia nervosa: an exploratory neuroimaging study

**DOI:** 10.1101/497669

**Authors:** Amy E. Miles, Allan S. Kaplan, Leon French, Aristotle N. Voineskos

## Abstract

Anorexia nervosa (AN) is a highly heritable psychiatric disorder characterized by starvation and emaciation and associated with changes in brain structure. The precise nature of these changes remains unclear, as does their developmental time course and capacity for reversal with weight restoration. In this exploratory neuroimaging study, we sought to characterize changes in white matter microstructure in women with acute and remitted AN. Diffusion-weighted MRI data was collected from underweight women with a current diagnosis of AN (acAN: n=23), weight-recovered women with a past diagnosis of AN (recAN: n=23), and age-matched healthy control women (HC: n=24). Image processing and analysis were performed with Tract-Based Spatial Statistics, part of FSL, and group differences in voxel-wise, brain-wide fractional anisotropy (FA) and mean diffusivity (MD), indices of white matter microstructure, were tested with nonparametric permutation and threshold free cluster enhancement. No significant main effect of group on FA was identified. A significant main effect of group on MD was observed in a large cluster covering 9.2% of white matter and including substantial portions of the corpus callosum, corona radiata, internal capsule, and superior longitudinal fasciculus, and post hoc analyses revealed similar effects of group on axial diffusivity (AD) and radial diffusivity (RD). Cluster-wise MD was significantly higher in acAN participants (+3.8%) and recAN participants (+2.9%) than healthy controls, and the same was true for cluster-wise AD and RD. Trait-based increases in diffusivity, consistent with atypical myelination and impaired axon integrity, suggest a link between altered white matter microstructure and vulnerability to AN, and evidence of reduced oligodendrocyte density in AN provides further support for this hypothesis. Potential mechanisms of action include atypical neurodevelopment and systemic inflammation.

## INTRODUCTION

Anorexia nervosa (AN) is a complex psychiatric disorder characterized by persistent restriction of energy leading to severely low body weight (for a review, see Treasure et al., 2015). AN typically emerges during adolescence, and it disproportionately impacts women. In the absence of evidence-based treatment, AN prognosis is relatively poor; mortality is estimated at ten percent, and fewer than half of patients achieve full recovery.

Despite strong evidence of heritability, consistent identification of neurodevelopmental risk factors (e.g. perinatal complications, childhood trauma), and robust documentation of pathophysiological consequences (e.g. fluid and electrolyte imbalance, endocrine dysregulation) (Treasure et al., 2015), investigation into the neurobiological correlates of AN has been limited. Moreover, it has focused primarily on changes in grey matter morphology (for a review, see Seitz, Herpertz-Dahlmann, & Konrad, 2016), with less attention paid to variations in white matter microstructure.

White matter tracts are densely-packed bundles of myelinated axons, neural projections that carry information between cells. These tracts develop over childhood and adolescence as myelinating glial cells (i.e. oligodendrocytes) surround axons, providing protection and electrical insulation and optimizing signal transmission, and their microstructure is moderately heritable (Voineskos, 2015). As such, additive genetic factors and early life stress exposure could drive atypical white matter development, and subsequent disruptions in coordinated information flow could confer vulnerability to AN. Likewise, starvation could give rise to white matter degeneration or disorganization that further hinders complex processing and contributes to disease chronicity.

Properties of white matter microstructure can be inferred from indices of diffusion derived from diffusion tensor imaging (DTI), an MRI method that uses patterns of water displacement to model tissue properties (Alexander, Lee, Lazar, & Field, 2007). These indices include fractional anisotropy (FA), a measure of directional-dependence of diffusion with higher values indicating greater fiber integrity, and mean diffusivity (MD), a measure of overall magnitude of diffusion with lower values indicating greater fiber integrity.

Several studies of DTI in AN have been published (for a review, see Monzon, Hay, Foroughi, & Touyz, 2016), and each has reported white matter impairment. However, all have been limited in scope; none has assessed microstructure in women with acute and remitted AN, and most have measured FA exclusively. As such, the primary aim of this study is to identify and characterize state-dependent and trait-based variations in white matter microstructure by comparing FA and MD in women with acute and remitted AN and healthy controls.

Including women with acute and remitted AN is paramount as it facilitates discrimination between variations in microstructure that are unique to symptomatic patients (i.e. state-dependent) and variations in microstructure that are independent of starvation state (i.e. trait-based) (Walsh & Devlin, 1998). This distinction has important implications for prevention and intervention as the former could intensify treatment resistance during acute illness while the latter could facilitate onset during critical periods in development or increase susceptibility to relapse after weight restoration. Likewise, testing group differences in FA and MD is imperative as the former is a highly sensitive but non-specific marker of white matter microstructure that provides limited information about neuropathology when considered in isolation (Alexander et al., 2007).

Although this study is exploratory, we have several hypotheses. First, we expect to observe state-dependent decreases in FA and state-dependent increases in MD in the fornix and superior longitudinal fasciculus (SLF), white matter tracts implicated in affective and body-centric processing, respectively, deficits in which are particularly pronounced during acute illness (Catani & Thiebautdeschotten, 2008; Treasure et al., 2015). Second, we expect to observe trait-based decreases in FA and trait-based increases in MD in the cingulum and anterior internal capsule/corona radiata, white matter tracts implicated in cognitive control and reward and reinforcement, respectively, deficits in which predate diagnosis and persist after weight recovery (Catani & Thiebautdeschotten, 2008; Treasure et al., 2015).

## METHODS AND MATERIALS

### Sample Recruitment

The current sample included 70 volunteers: 23 women with acute AN (acAN: 18 – 47 years), 23 women with remitted AN (recAN: 19 – 43 years), and 24 age-matched healthy control women (HC: 21 – 47 years).

Participants with a lifetime diagnosis of AN were recruited from hospital and community-based clinics in the Greater Toronto Area. Participants assigned to the acAN group met the following criteria: [1] female, aged 18 – 48; [2] current AN diagnosis; [3] BMI < 18.0 kg/m^2^. Participants assigned to the recAN group met the following criteria: [1] female, aged 18 – 48; [2] past AN diagnosis; [3] BMI > 18.5 kg/m^2^, as maintained for twelve consecutive months; [4] abstinence from bingeing, purging, and restrictive eating, as maintained for twelve consecutive months. Participants were excluded from both groups if they had a history of psychosis or met criteria for a substance use disorder.

Healthy controls were selected from a database maintained by the co-PI, Dr. Aristotle Voineskos, based on the following criteria: [1] female, aged 18 – 48; [2] BMI > 18.5 kg/m^2^ and BMI < 30.0 kg/m^2^. Subjects in this database were recruited as part of an ongoing neuroimaging study for which they were interviewed by a psychiatrist, assessed and weighed by research staff, and scanned in accordance with the protocol described below. A query of this database identified twenty-four subjects who met eligibility criteria, and all were included. As per project guidelines, these subjects had no lifetime psychiatric history and no family history of psychosis, and they did not meet criteria for a current substance use disorder or have a positive urine toxicology screen. They were assessed and imaged between February 2015 and July 2017.

Participants were excluded from all groups if they were pregnant or otherwise medically unfit for imaging or if they had a history of severe head trauma or neurological disorder.

The study was approved by the Centre for Addiction and Mental Health Research Ethics Board, and all participants gave written informed consent.

### Clinical Assessment

A comprehensive clinical battery was used to assess participants with a lifetime diagnosis of AN. As part of this battery: [1] a semi-structured interview, administered by the first author, was used to collect demographic and clinical information; [2] the Structured Clinical Interview for Diagnosis, Research Version (SCID-I/P: First, Spitzer, Gibbon, & Williams, 2002), as modified to reflect guidelines from the Diagnostic and Statistical Manual of Mental Disorders, Fifth Edition (American Psychiatric Association, 2013), was used to confirm AN diagnosis, screen for psychosis and substance use, and report comorbid anxiety and mood disorders; [3] the Eating Disorder Examination Questionnaire (EDE-Q 6.0: Fairburn & Beglin, 2008) was used to characterize disease presentation.

### Image Acquisition and Processing

MRI data was acquired on a 3T scanner (Echospeed; General Electric Medical Systems). For DTI, sixty diffusion-weighted (b = 1000 s/mm^2^) and five baseline scans (b = 0 s/mm^2^) were acquired using an echo planar imaging sequence with dual spin-echo and the following parameters: TE = 88 ms; TR = 8800ms; FOV = 256 mm; 128x128 encoding steps; 2.0mm isotropic voxels, no gap. For anatomical co-registration, high-resolution 3D T1-weighted structural scans were acquired using a fast-spoiled gradient-recalled echo sequence with the following parameters: TE = 3ms; TR = 8.2ms; TI = 650ms; α = 8°; FOV = 24cm; NEX = 1; 0.9mm isotropic voxels, no gap.

MRI preprocessing was carried out with tools from the FMRIB Software Library (FSL) (https://fsl.fmrib.ox.ac.uk, version 5.0.10) (Smith et al., 2004). Steps included: [1] correcting for head motion and distortion induced by eddy currents; [2] fitting tensors at each voxel; [3] removing non-brain tissue.

DTI analysis was carried out with Tract-Based Spatial Statistics (TBSS) (Smith et al., 2006), also part of FSL. This method involves: [1] aligning subjects’ raw FA data to the FMRIB58_FA template using non-linear registration; [2] averaging subjects’ raw FA data to create a mean FA image; [3] thinning the mean FA image to create a mean FA skeleton representing the center of tracts common to all participants; [4] thresholding the mean FA skeleton to exclude non-white matter and regions of high individual variability; [5] projecting subjects’ normalized FA data onto the mean FA skeleton; [6] running voxel-wise cross-subject statistics on skeletonized data, as described below.

As part of a comprehensive quality control procedure, raw diffusion data, native-space FA images, and standard-space FA images were visually inspected by a trained examiner, E.D.

Whole-brain tissue volumes were estimated from T1-weighted images using the Freesurfer volume-based processing pipeline. This method has been described in detail elsewhere (Fischl et al., 2002, 2004), and its implementation is summarized in Miles, Voineskos, French, & Kaplan (2018).

### Primary Statistical Analyses

Statistical analyses were performed using RStudio (RStudio Team, version 0.99.489) and FSL *Randomise* (Winkler, Ridgway, Webster, Smith, & Nichols, 2014).

For analyses of demographic and clinical characteristics, main effect of group was tested with univariate analyses of variance or covariance (ANCOVA) and chi-squared tests, as appropriate, and post hoc pair-wise comparisons were tested with Tukey HSD or their significance values were adjusted using the Holm-Bonferroni method.

For analyses of white matter microstructure, main effect of group was modeled with age as a nuisance variable and tested with nonparametric permutation-based inference (10,000 permutations) and threshold-free cluster enhancement (Smith & Nichols, 2009). Voxel-wise, brain-wide analyses were repeated for FA and MD and clusters were identified using a family-wise error-corrected threshold (p_FWER_ < .025) and labeled according to the JHU ICBM-DTI-81 white matter atlas (Mori et al., 2008).

### Post Hoc Statistical Analyses

FSL command-line utilities were used to extract cluster-wise statistics for post hoc analysis.

Main effect of group on cluster-wise FA and MD was modeled with age as a nuisance variable and confirmed with univariate ANCOVA, and group differences therein were tested with Tukey HSD.

Cluster-wise analyses were repeated for axial diffusivity (AD) and radial diffusivity (RD), additional DTI-derived measures that index diffusion parallel and perpendicular to the primary axis (i.e. white matter fiber bundle) and are thought to reflect axon and myelin integrity, respectively (Alexander, Lee, Lazar, & Field, 2007). To account for testing of multiple DTI measures, a Bonferroni-adjusted threshold was used to determine significance of F statistics.

Multiple regression was used to assess the impact of the following clinical variables on cluster-wise FA and MD: [1] current treatment with one or more selective serotonin reuptake inhibitors (SSRIs) or atypical antipsychotics (AAPs); [2] lifetime diagnosis of major depression (MDD); [3] lifetime diagnosis of generalized anxiety disorder (GAD), obsessive-compulsive disorder (OCD), panic disorder (PD), or post-traumatic stress disorder (PTSD).

To account for testing of multiple predictor values, a Bonferroni-adjusted threshold was used to determine significance of t-statistics.

## RESULTS

Demographic and clinical variables and DTI measures are summarized in Tables 1 and 2, respectively, and predictors of white matter microstructure are summarized in Table 3. Clusters of significant between-groups difference in FA and MD are depicted in Figure 1.

**FIGURE 1.**
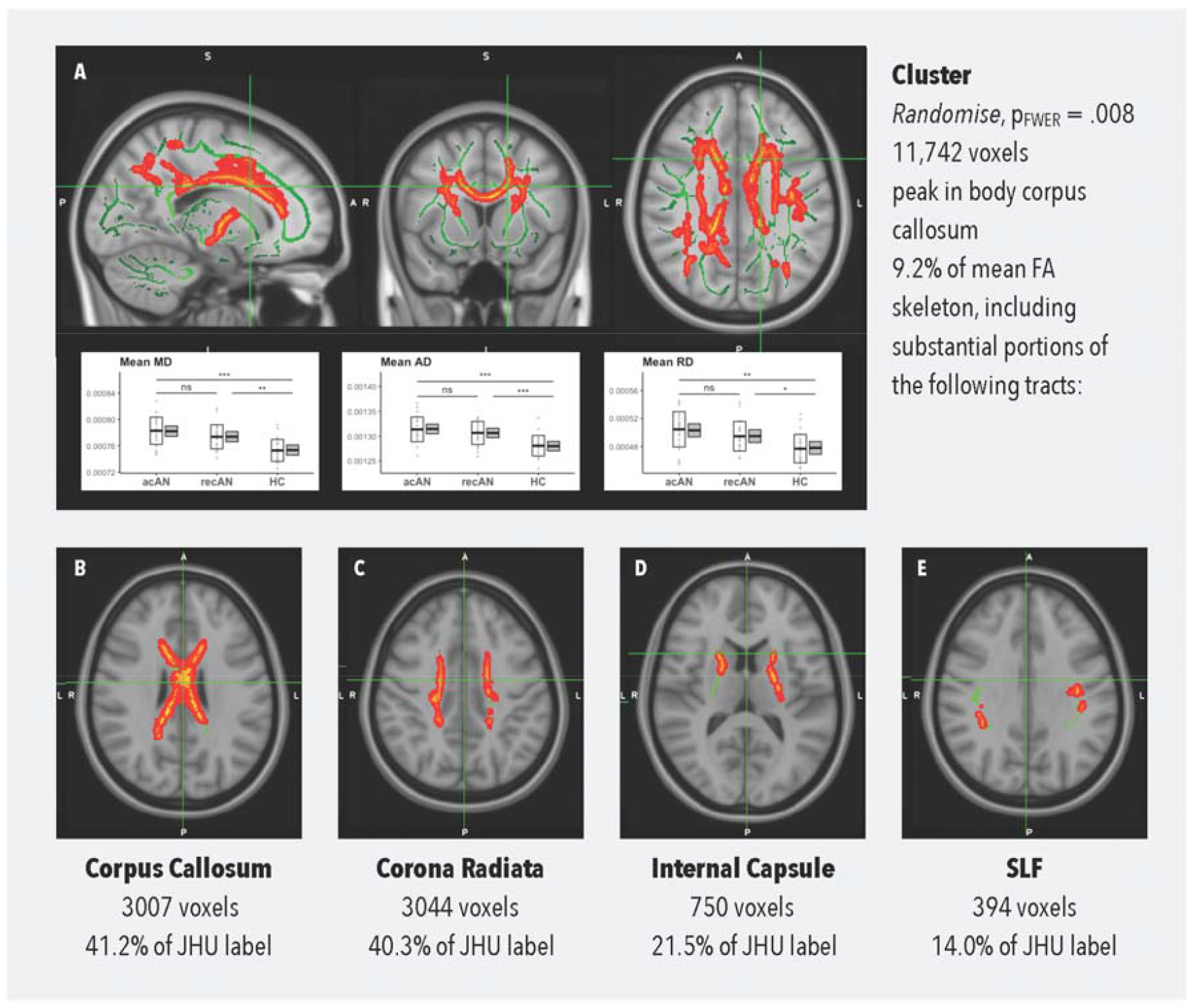
ANCOVA RESULTS: Cluster in which there is a significant main effect of group on peak MD, as overlaid onto the mean FA skeleton (A) and masked by JHU label (B, C, D, E). p_FWER_ < 0.05 (*), 0.01 (**), 0.001 (***); acAN = acute anorexia; AD = axial diffusivity; FA = fractional anisotropy; FWER = family-wise error-corrected; HC = healthy control; JHU = JHU ICBM-DTI-81 white matter atlas; MD = mean diffusivity; ns = not significant; RD = radial diffusivity; recAN = remitted anorexia; SLF = superior longitudinal fasciculus

**TABLE 1.** SAMPLE CHARACTERISTICS: Demographic and clinical characteristics and group differences therein. **p_FWER_ < 0.05, *p_FWER_ < 0.10***; AAP = atypical antipsychotic; acAN = acute anorexia; BMI = body mass index; EDE-Q = Eating Disorder Examination Questionnaire, Version 6.0; FWER = family-wise error-corrected; GAD = generalized anxiety disorder; GLM = general linear model; HC = healthy control; MDD = major depressive disorder; n/a = not applicable; OCD = obsessive-compulsive disorder; PD = panic disorder; PTSD = post-traumatic stress disorder; recAN = remitted anorexia; SD = standard deviation; SSRI = selective serotonin reuptake inhibitor

|  | Mean (SD) and cell counts |  |  | GLM and chi-squared test statistics |  |  |  |
| --- | --- | --- | --- | --- | --- | --- | --- |
|  | acAN<sub>(1)</sub> | recAN<sub>(2)</sub> | HC<sub>(3)</sub> | F | T<sub>(1,2)</sub> | T<sub>(1,3)</sub> | T<sub>(2,3)</sub> |
| **N** | 23 | 23 | 24 | n/a | n/a | n/a | n/a |
| **Age**, years | 29.6 (8.2) | 26.7 (6.2) | 25.0 (5.7) | 2.71 | n/a | n/a | n/a |
| **BMI**, kg/m<sup>2</sup> | 16.0 (1.4) | 20.9 (1.8) | 22.3 (2.2) | **75.65** | **9.08** | **11.75** | **2.57** |
| **Education**, years, age-adjusted | 14.8 (0.3) | 15.7 (0.3) | 16.4 (0.3) | **5.60** | 2.00 | **3.34** | 1.41 |
| **Age at Onset**, years | 15.0 (5.1) | 15.0 (3.0) | n/a | n/a | 0.04 | n/a | n/a |
| **Illness Duration**, years, age-adjusted | 12.4 (1.0) | 10.1 (1.0) | n/a | n/a | **12.15** | n/a | n/a |
| **EDE-Q Risk Composite** | 4.0 (1.3) | 2.3 (1.7) | n/a | n/a | **3.71** | n/a | n/a |
| **Race**, white:non-white | 21:2 | 18:5 | 14:10 | **7.06** | 0.67 | **5.09** | 1.33 |
| **Handedness**, right:left | 22:1 | 19:4 | 20:4 | 2.22 | n/a | n/a | n/a |
| **Subtype**, restricting:purging | 8:15 | 16:7 | n/a | n/a | **4.27** | n/a | n/a |
| **Current AAP or SSRI**, yes:no | 16:7 | 9:14 | 0:24 | **24.93** | **3.15** | **22.31** | **9.23** |
| **Lifetime MDD**, yes:no | 19:4 | 9:14 | 0:24 | **33.41** | **7.39** | **29.94** | **9.23** |
| **Lifetime GAD, OCD, PD, or PTSD**, yes:no | 15:8 | 15:8 | 0:24 | **27.39** | 0.00 | **20.09** | **20.09** |

**TABLE 2.** REGIONS OF INTEREST: Cluster and tract means and group differences therein. **p_FWER_** < 0.05, *p_FWER_ < 0.10*; acAN = acute anorexia; AD = axial diffusivity; FWER = family-wise error-corrected; GLM = general linear model; HC = healthy control; MD = mean diffusivity; n/a = not applicable; RD = radial diffusivity; recAN = remitted anorexia; SLF = superior longitudinal fasciculus

|  | Multiple regression T statistics |  |  |  |  |  |
| --- | --- | --- | --- | --- | --- | --- |
|  | ANX | MDD | Rx | Edu. | AN-BP | Dur. |
| **Cluster, MD** | -0.55 | **-2.55** | 0.13 | 1.30 | -1.59 | 1.22 |
| **Corpus Callosum, MD** | -1.23 | **-3.38** | 0.13 | 0.91 | -2.31 | 1.31 |
| **Corona Radiata, MD** | -0.12 | -2.37 | 0.11 | 1.27 | -1.07 | 0.59 |
| **Internal Capsule, MD** | 0.07 | -1.68 | -0.17 | 1.51 | -1.35 | -0.93 |
| **SLF, MD** | -0.15 | -1.16 | -0.18 | 1.01 | -0.68 | 0.33 |

**TABLE 3.**
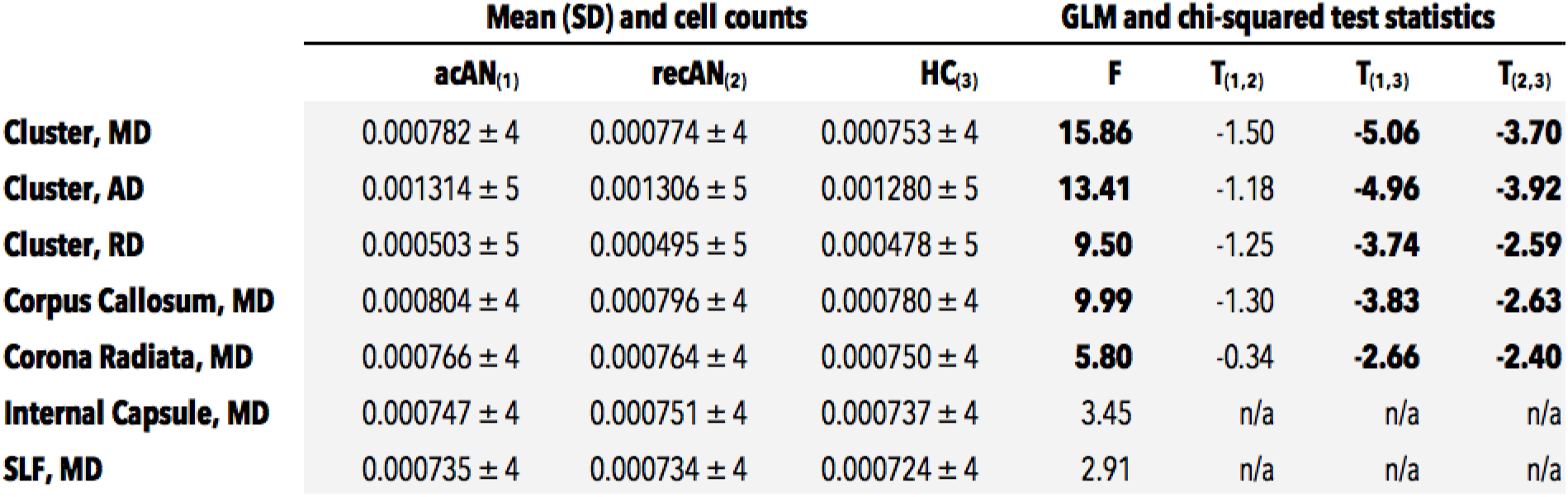
CLINICAL PREDICTORS OF MICROSTRUCTURE: Effects of clinical variables of interest on white matter microstructure, controlling for group and age. **p_FWER_ < 0.05**; AN-BP = binge-purge subtype anorexia; ANX = lifetime diagnosis of generalized anxiety disorder, obsessive-compulsive disorder, panic disorder, or post-traumatic stress disorder; Dur. = illness duration; Edu. = education; FWER = family-wise error-corrected; MD = mean diffusivity; MDD = lifetime diagnosis of major depressive disorder; Rx = current treatment with one or more atypical antipsychotics or selective serotonin reuptake inhibitors; SLF = superior longitudinal fasciculus

|  | Mean (SD) and cell counts |  |  | GLM and chi-squared test statistics |  |  |  |
| --- | --- | --- | --- | --- | --- | --- | --- |
|  | acAN<sub>(1)</sub> | recAN<sub>(2)</sub> | HC<sub>(3)</sub> | F | T<sub>(1,2)</sub> | T<sub>(1,3)</sub> | T<sub>(2,3)</sub> |
| **Cluster, MD** | 0.000782 ± 4 | 0.000774 ± 4 | 0.000753 ± 4 | **15.86** | -1.50 | **-5.06** | **-3.70** |
| **Cluster, AD** | 0.001314 ± 5 | 0.001306 ± 5 | 0.001280 ± 5 | **13.41** | -1.18 | **-4.96** | **-3.92** |
| **Cluster, RD** | 0.000503 ± 5 | 0.000495 ± 5 | 0.000478 ± 5 | **9.50** | -1.25 | **-3.74** | **-2.59** |
| **Corpus Callosum, MD** | 0.000804 ± 4 | 0.000796 ± 4 | 0.000780 ± 4 | **9.99** | -1.30 | **-3.83** | **-2.63** |
| **Corona Radiata, MD** | 0.000766 ± 4 | 0.000764 ± 4 | 0.000750 ± 4 | **5.80** | -0.34 | **-2.66** | **-2.40** |
| **Internal Capsule, MD** | 0.000747 ± 4 | 0.000751 ± 4 | 0.000737 ± 4 | 3.45 | n/a | n/a | n/a |
| **SLF, MD** | 0.000735 ± 4 | 0.000734 ± 4 | 0.000724 ± 4 | 2.91 | n/a | n/a | n/a |

### Sample Characteristics

Mean age was comparable across groups as was number of right-handed participants. Mean education, as adjusted for age, was significantly lower in acAN participants than healthy controls (t = 3.335, p_FWER_ = .004), and there was a trend towards less racial diversity in the former group (χ^2^ = 5.093, p_FWER_ = .072).

As anticipated, mean BMI was significantly lower in acAN participants than recAN participants (t = 9.084, p_FWER_ < .001) and healthy controls (t = 11.747, p_FWER_ < 001), and rates of current psychotropic medication use, lifetime depression diagnosis, and lifetime anxiety-spectrum diagnosis were significantly higher in acAN participants and recAN participants than healthy controls (t > 9.225, p_FWER_ < .006).

As further anticipated, mean EDE-Q 6.0 composite score was significantly higher in acAN participants than recAN participants (t = 3.709, p = .001), and the same was true for age-adjusted illness duration (t = 12.150, p = .001) and rates of binge-purge subtype anorexia (χ^2^ = 4.269, p = .039), lifetime depression diagnosis (χ^2^ = 7.393, p_FWER_ = .007), and current psychotropic medication use (χ^2^ = 3.154, p_FWER_ = .076), the latter at trend level. Mean age at onset was comparable between groups, as was rate of lifetime anxiety-spectrum diagnosis.

### Group Differences in White Matter Microstructure

We identified no clusters in which there was a significant main effect of group on peak FA. We detected a significant main effect of group on peak MD (F = 18.061, p_FWER_ = .008) in one cluster covering 9.2% of the white matter skeleton and including substantial portions of the corpus callosum, corona radiata, internal capsule, and SLF, and post hoc analyses confirmed this result (MD: F = 15.859, p < .001) and revealed comparable effects of group on AD (F = 13.405, p_FWER_ < .001) and RD (F = 9.495, p_FWER_ < .001). In each case, cluster-wise diffusivity was highest in acAN participants and lowest in healthy controls, and acAN and recAN groups differed significantly from healthy controls but not from one another. Cluster-wise MD was 3.8% higher in acAN participants than healthy controls, and it was 2.9% higher in recAN participants than healthy controls.

Given the size and spatial extent of the aforementioned cluster, we tested main effect of group on mean MD in each of the major white matter tracts (i.e. corpus callosum, corona radiata, internal capsule, and SLF) represented therein, using a Bonferroni-adjusted threshold to determine significance of F statistics. In doing so, we detected a significant main effect of group on mean MD in the corpus callosum (F = 9.992, p_FWER_ = .001) and corona radiata (F = 5.802, p_FWER_ = .019), and we observed a consistent pattern of between-group difference whereby tract-wise MD was significantly higher in acAN and recAN participants than healthy controls.

### Clinical Predictors of White Matter Microstructure

Neither current use of SSRIs or AAPs nor lifetime diagnosis of GAD, OCD, PD, or PTSD was a significant predictor of cluster/tract-wise MD after accounting for effects of group and age and correcting for multiple comparisons. The same was true of education, disease subtype, and illness duration, as tested with additional regression analyses given significant group differences therein.

Lifetime depression diagnosis was not significantly related to cluster-wise MD, but it was a significant predictor of mean MD in the corpus callosum (t = −3.375, p_FWER_ = .008). Including lifetime depression diagnosis as an additional covariate had a negligible impact on main effect of group (cluster, F = 17.180, p < .001; corpus callosum, F = 11.564, p < .001).

Neither BMI nor EDE-Q composite score was significantly associated with cluster-wise MD in acAN participants (BMI: r = -0.037, p = 0.868; EDE-Q: r = -0.194, p = 0.376) or recAN participants (BMI: r = -0.079, p = 0.719; EDE-Q: r = -0.112, p = 0.611).

## DISCUSSION

In this DTI study in women with and without AN, we found no evidence of state-dependent changes in white matter microstructure.

### Trait-Based Increases in Mean Diffusivity

In contrast, we detected trait-based variations in microstructure (MD, AD, RD: acAN, recAN > HC) in a large cluster covering substantial portions of the corpus callosum, corona radiata, internal capsule, and SLF, and the same was true when averaged over each of the largest tracts therein (i.e. corpus callosum, corona radiata). This finding is somewhat consistent with previous studies of white matter in adult AN; of seven DTI studies published to date, four have reported altered microstructure in at least one of the aforementioned tracts (Hayes et al., 2015; Shott, Pryor, Yang, & Frank, 2016; Via et al., 2014; Yau et al., 2013), although none has reported increased MD.

Site-specific microstructural impairment could have significant bearing on AN psychopathology. Altered corpus callosum microstructure could give rise to altered taste processing, selective attention to disorder-salient stimuli, and difficulty identifying emotional state and interpreting behavior, prominent features of AN (Treasure, Zipfel, et al., 2015) and complex processes that require interhemispheric integration, by disrupting information flow between homologous regions in left and right hemispheres (Catani & Thiebautdeschotten, 2008; Mori et al., 2008). Likewise, altered internal capsule/corona radiata microstructure could give rise to excessive behavioral avoidance and impaired response inhibition, putative endophenotypes of AN (Treasure, Zipfel, et al., 2015), by disrupting information flow among cortical and subcortical regions implicated in reward and reinforcement and goal-directed behavior (i.e. frontoparietal cortex, striatum) (Catani & Thiebautdeschotten, 2008; Mori et al., 2008). Finally, altered SLF microstructure could give rise to body size misperception, a hallmark of AN and one of its most persistent symptoms (Cash & Deagle, 1997; Fairburn, Cooper, & Shafran, 2003), by disrupting information flow among non-adjacent cortical regions (i.e. frontal and opercular cortex and inferior parietal, superior parietal, and superior temporal cortex) implicated in visual attention, spatial perception, and body-specific processing (Catani & Thiebautdeschotten, 2008; Makris et al., 2005; Mori et al., 2008).

Moreover, site-specific microstructural impairment could emerge early in development and confer vulnerability to disease. Comparably large increases in MD in women with acute and remitted AN supports this hypothesis, as does the absence of association between microstructural impairment and illness duration. Furthermore, high estimates of heritability in regions of interest (h^2^ = 0.86 – 0.90) (Kochunov et al., 2015) and comparable findings in animal models of abuse and neglect (Teicher et al., 2003), developmental risk factors for AN (Favaro et al., 2010), and in patients with autism spectrum disorder (Alexander et al., 2007), a neurodevelopmental disorder, symptoms of which are overrepresented in AN (Westwood, Mandy, & Tchanturia, 2017), suggest premorbid ontogeny.

Together, trait-based increases in MD, AD, and RD point towards de/dysmyelination and axonal injury as potential mechanisms of action (Alexander et al., 2007; Della Nave et al., 2011; Song et al., 2002, 2005). Myelination occurs over the course childhood and early adolescence as glial cells proliferate and encapsulate axons, producing a characteristic reduction in RD as diffusion become more directionally-dependent (Lebel & Beaulieu, 2011). As such, insufficient production or pathological loss of myelin could drive observed increases in RD. Evidence of de/dysmyelination in AN comes from post-mortem analysis of gene expression profiles from the dorsolateral prefrontal cortex. These profiles were collected by Jaffe et al. (2014) for a study comparing gene expression in 31 patients with obsessive psychiatric syndromes and 102 healthy controls (GSE60190). A filtered version of this dataset was obtained from the Gemma database (Zoubarev et al., 2012) and expression profiles from five AN cases and eight matched controls were used to estimate relative abundance of astrocytes, neurons, oligodendrocytes, oligodendrocyte precursors, microglia and endothelial cells (Darmanis et al., 2015; Mancarci et al., 2017). Comparison thereof revealed significant underrepresentation of oligodendrocytes in the AN group (t = -4.59, p_FWER_ = 0.008), and direct testing of thirty probes, corresponding to sixteen myelination-related genes selected from a set identified by Chavarria-Siles et al. (2016), revealed significantly lower expression of *KLK6, MOG,* and *TF* in cases than controls (p_FWER_ = .047).

Axonal injury is a consequence of physical trauma, infection, and inflammation associated with increased membrane permeability and cellular degeneration and characterized by changes in AD (Medana & Esiri, 2003; Moreno et al., 2011). As such, increased tissue water content due to serious injury or chronic illness could drive observed increases in AD. The reported link between AN risk and family history of auto-immune or auto-inflammatory disorders provides preliminary support for this hypothesis (Zerwas et al., 2017).

Further research is crucial as our findings only partially replicate those of other groups. Increased sensitivity due to greater scanner strength (3T vs 1.5T) and better tensor fitting due to more gradient directions (60 vs 25) could account for these findings, as could larger group sizes, especially in women with remitted AN (23 vs 12). The same is true of less CSF contamination, an artifact to which the fornix is particularly susceptible (Berlot, Metzler-Baddeley, Jones, & O’Sullivan, 2014), and less partial voluming due to atrophy; we detected no significant group differences in CSF volume (acAN vs. HC, t = -1.071, p_FWER_ = 0.535; recAN vs. HC, t = 1.462, p_FWER_ = 0.316), grey matter volume (acAN vs. HC, t = 1.374, p_FWER_ = 0.360; recAN vs. HC, t = – 1.408, p_FWER_ = 0.343), or white matter volume (acAN vs. HC, t = 0.510, p_FWER_ = 0.867; recAN vs. HC, t = -1.700, p_FWER_ = 0.213).

Further research is also needed to explore clinical implications and test hypothesized mechanisms, and the possibility of competing effects should be carefully considered. This is particularly true given observed increases in RD (associated with lower FA) and increases in AD (associated with higher FA) that suggest cooccurring de/dysmyelination and axonal injury and could account for non-significant group differences in FA.

### Limitations and Future Directions

This exploratory neuroimaging study is limited by its relatively small sample size and cross-sectional design, and inference, made with limited empirical support for hypothesized mechanisms, is hampered by sample composition. Particularly notable shortcomings include: [1] selection of healthy controls from an existing database, precluding pair-wise age-matching and uniform assessment; [2] assignment to acAN and recAN groups based on BMI, not BMI and EDE-Q score; [3] significant group differences in disease subtype and illness duration. The latter shortcoming is particularly important as it threatens the underlying assumption that acAN and recAN participants are sampled from the same clinical population.

Despite these weaknesses, this study makes several important contributions to our understanding of AN neurobiology. First, it replicates previous reports of altered white matter microstructure in women with acute and remitted AN, identifying trait-based increases in MD in major association, commissural, and projection tracts. Second, it implicates de/dysmyelination and axonal injury in the pathophysiology of AN. Finally, it points towards altered neurodevelopmental patterning and systemic inflammation in the pathogenesis of AN. Replication in a larger sample, parsed by subtype and tracked longitudinally, could provide strong validation of these findings, as could replication in healthy siblings.

## ACKNOWLEDGEMENTS

We thank Andrew Jaffe and Joel E. Kleinman for providing eating disorder subtype information for the GSE60190 expression dataset.

We also thank staff at the CAMH Research Imaging Centre and members of the Kimel Family Translational Imaging-Genetics Laboratory, particularly Dr. Erin Dickie, for their assistance with MRI acquisition and processing.

## COMPLIANCE WITH ETHICAL STANDARDS

### Funding

This study was funded by the AFP Innovation Fund (CAM-114-001, CAM-116-003).

### Ethical Approval

All procedures performed in studies involving human participants were in accordance with the ethical standards of the institutional and/or national research committee and with the 1964 Helsinki declaration and its later amendments or comparable ethical standards.

This article does not contain any studies with animals performed by any of the authors.

### Informed Consent

Informed consent was obtained from all individual participants included in the study.

